# PIB: Parallel ion beam etching of sections collected on wafer for ultra large-scale connectomics

**DOI:** 10.1101/2025.04.25.650569

**Authors:** Hongtu Ma, Yanan Lv, Minggang Wang, Zhuangzhuang Zhao, Jing Liu, Xiaohui Dong, Yanchao Zhang, Jinyue Guo, Bohao Chen, Lina Zhang, Sheng Chang, Haoran Chen, Hao Zhai, Linlin Li, Xi Chen, Hua Han

## Abstract

We developed a parallel ion beam etching device that simultaneously thins hundreds of sections on a whole 4-inch wafer at 20 nm steps, enabling sub-centimeter volumetric imaging with high axial resolution. Compared with traditional volume electron microscopy, PIB preserves in-situ continuity within sections and simplifies inter-section registration. Its high axial resolution facilitates reliable neuron segmentation and tracing with reduced proofreading, providing an efficient route toward ultra-large-scale connectomics, such as whole-mouse-brain reconstruction.

## (71 字)Introduction

High-resolution volume electron microscopy (vEM) imaging of biological samples is critical in connectomics, where the goal is to map neural circuits in unprecedented detail [1–3]. As this field progresses, efforts to achieve high-resolution imaging of large-scale brain samples, have become a priority for future connectomics research [4]. Traditional vEM methods for large specimens rely heavily on ultramicrotomes to produce thousands of ultrathin serial sections [5–8]. However, these methods are limited in terms of achievable axial resolution, as the precision of section thickness is restricted by the mechanical limits of ultramicrotomes, particularly in maintaining stable operation over extended periods [9]. Additionally, the ultrathin sectioning process is highly sensitive to environmental factors such as vibrations and temperature fluctuations, which frequently result in sectioning failures and lower throughput [10]. Consequently, Section-based vEM methods typically have an axial resolution above 30 nm and suffer from low imaging efficiency, hindering the detailed analysis of neuronal connections, reducing the accuracy of automatic 3D segmentation, and increasing the burden of manual proofreading. In contrast to the limited processing area of Focused Ion Beam Scanning Electron Microscopy (FIB-SEM), Gas Cluster Ion Beam Scanning Electron Microscopy (GCIB-SEM) can achieve high axial resolution while thinning samples over a 7.5×7.5 mm^2^ area at one operation process [11, 12]. This capability allows for the processing of larger samples, yet for whole mouse brains, the inability to efficiently thin several sections in parallel means that the required throughput cannot be achieved. To address these challenges, we develop a section-thinning device that employs parallel ion beam etching (PIB) technology, adapted from the semiconductor field. By reducing incident energy of the ion beam, the device can effectively process sections collected on a whole 4-inch wafer simultaneously with 20nm thinning step. This advancement addresses the axial resolution requirements for ultra large brain samples and significantly enhances the efficiency, scalability and robustness of sample preparation in connectomics, facilitating the study of larger brain regions with higher accuracy and reliability.

In GCIB-SEM, the beam diameter is approximately 250 µm, which necessitates the use of an electrostatic scanning plate to enable rapid deflection of the ion beam across the sample surface [12]. This limitation constrains the effective processing area to less than 10 mm. Moreover, to minimize thinning artifacts, the ion beam was at a 30-degree glancing angle to the sample surface, which can lead to uneven thinning across larger samples. This non-uniformity needs additional post-processing steps to achieve the flatness of the imaged 3D volume. In contrast, our method generates a parallel ion beam that delivers uniform energy across a much larger area of up to 150 mm. This capability enables consistent thinning of hundreds of sections collected on a 4-inch wafer simultaneously, which is desirable for high-throughput electron microscopy imaging. Furthermore, the PIB method employs a shallower glancing angle of 5 degrees, significantly mitigating the impact of differential sputtering efficiency associated with varying atomic numbers in the sample. This design feature contributes to a more uniform thinning process, enhancing the overall quality and consistency of sample preparation in connectomics research.

Our PIB prototype device includes control system, vacuum system, RF power supply, ion source, neutralizer, and thinning chamber (Fig. 1a, Supplementary Fig. 1a, b, c). In contrast to ion guns such as FIB and GCIB, PIB generates a parallel ion beam with a diameter of up to 150 mm. This capability is enabled by the triple-grid regulation mechanism of the ion source (Fig. 1a, b, c, d; Supplementary Fig. 1d). In this design, the acceleration and ground grids primarily collimate the ion beam without changing its kinetic energy, while the beam energy is determined by the voltage applied to the screen grid (Supplementary Fig. 1d). The neutralizer emits low-energy electrons to compensate for the positive charges within the ion beam, thereby suppressing beam divergence and preventing charge accumulation on the sample surface. The relatively low energy and parallelism of the ion beam are essential for achieving uniform, large-area 20nm thinning of biological sections at one time (see *Methods* for working principle of ion source).

**Fig. 1.**
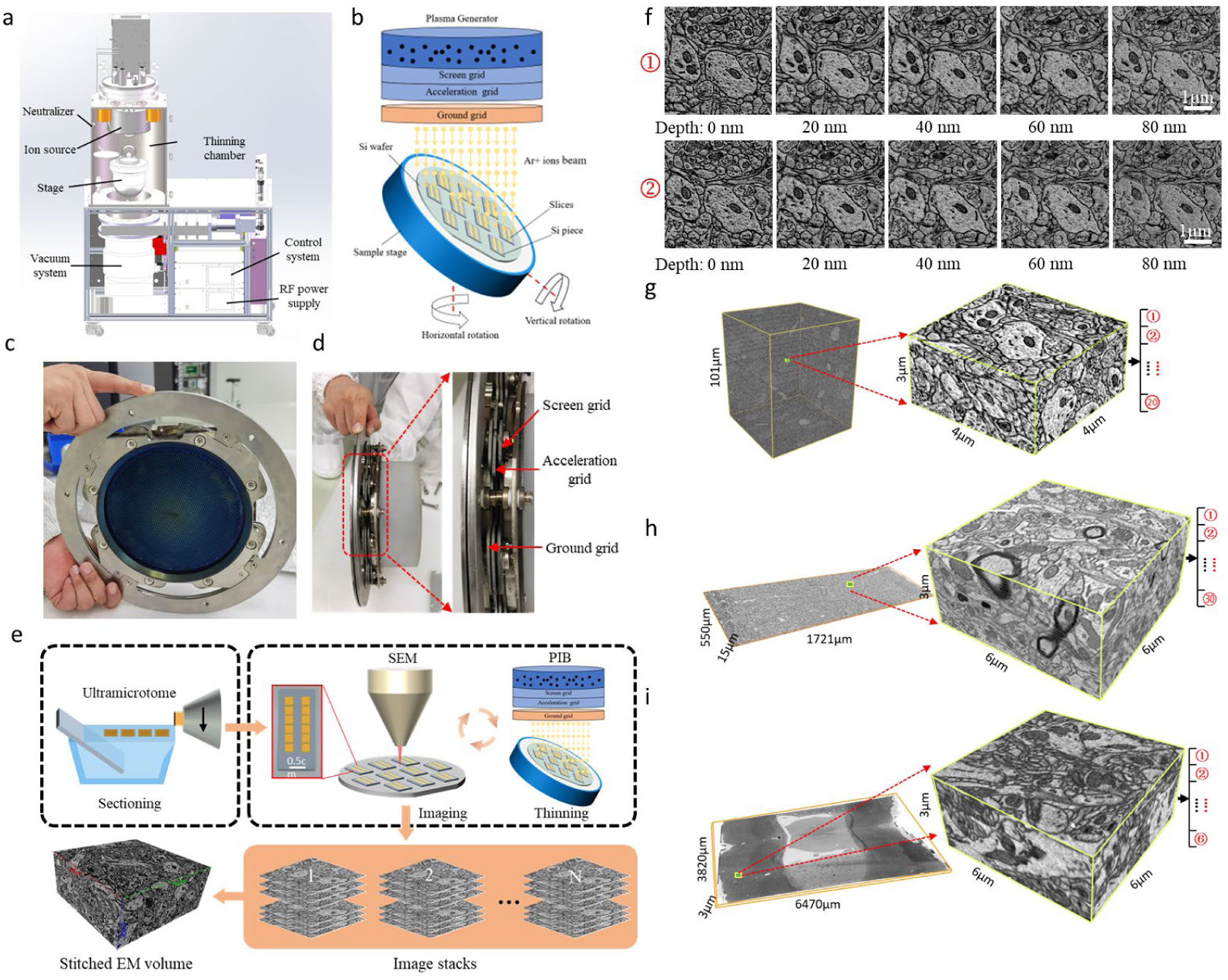
PIB-SEM overview. **a,** 3D model and actual appearance of the PIB prototype. **b**, Schematic diagram of PIB-SEM working principle. After the plasma is generated, it is extracted and shaped by the three-grid structure to form a large-area parallel ion beam for thinning sections. **c**, Front view of the ion source. **d,** Side view of the ion source showing the triple-grid structure, which includes the screen grid, acceleration grid, and ground grid. **e,** The workflow of PIB-SEM. First, the biological sample embedded in resin block are sliced using an ultramicrotome, and the numbered sections are collected onto silicon pieces. Next, all silicon pieces are placed into SEM for serial section imaging. After imaging, the silicon pieces are placed into the thinning chamber of PIB for section thinning, completing an PIB-SEM cycle. With several PIB-SEM cycles, we obtain the image stacks for all sections. Finally, the image stacks are aligned to form a three-dimensional volume of the biological sample. **f,** SEM images of two consecutive sections in the sub-volume at five different thinning depths. **g**, PIB-SEM 3D volume (83 μm × 83 μm × 101 μm) spanning 1018 sequential 100 nm sections of rat cortex, and a sub-volume (4 μm × 4 μm × 2 μm) spanning 20 sequential 100 nm sections (numbers denote individual thick sections). **h**, PIB-SEM 3D volume (1721 μm × 550 μm × 15 μm) spanning 150 sequential 100 nm sections from all layers of the rat cortex, and a sub-volume (6 μm × 6 μm × 3 μm) spanning 30 sequential 100 nm sections (numbers denote individual thick sections). **i**, PIB-SEM 3D volume (6470 μm × 3820 μm × 3 μm) spanning 6 sequential 500 nm sections of the rat brain, and a sub-volume (6 μm × 6 μm × 3 μm) spanning 6 sequential 500 nm sections (numbers denote individual thick sections).

The workflow of PIB-SEM is illustrated in Fig. 1e. The sample is cut into serial sections with a thickness of 100 nm or above using ultramicrotome. Compared to 30 nm sections, it is much easier to obtain qualified continuous sections at thicknesses ≥100 nm, which are free of distortion, such as wrinkle and tear. The ribbons of serial sections are then collected on hydrophilized silicon (Si) pieces. After imaging the region of interest of serial sections in SEM, these Si pieces are mounted in PIB to remove the surface of the sections with a thickness of 20 nm. Through multiple imaging and thinning cycles, we obtain EM images of all sections with a 20 nm axial resolution. Finally, the obtained image stack is aligned to generate the 3D volume of the sample.

## Results

To evaluate the performance of PIB-SEM for high-resolution volumetric imaging, we use the sample from prefrontal cortex of a rat (see *Methods* for prefrontal cortex of rat). A total of 1018 consecutive sections with 100 nm thickness are obtained and collected on three hydrophilized Si pieces (Supplementary Fig. 3b). After imaging in Zeiss Gemini SEM 300 with a BSD detector, these Si pieces are individually placed into the thinning chamber of PIB to remove the surface of the sections, which take about 12 min to thinning 20nm. Due to mechanical deviations of the ultramicrotome, we perform multiple cycles of imaging and thinning, with the number of cycles depending on the thickness of the thickest section. For those thinner sections, the images obtained in the last few cycles have no meaningful content, so we discard them in serial sections alignment (Supplementary Fig. 3). Two consecutive sections with 5 different thinning cycles are shown in Fig. 1f, where the continuity of neural substructures within and between sections are preserved. All sections are stitched into a volume of 83 μm × 83 μm × 101 μm. A sub-volume of 4 μm × 4 μm × 2 μm spanning 20 sections (Fig. 1g) and a sub-volume of 8 μm × 8 μm × 4 μm spanning 40 sections (Supplementary Fig. 4, Supplementary Video 1) are presented in detail.

To objectively evaluate the axial continuity of the PIB-SEM volume, we select the automated tape collection ultramicrotome SEM (ATUM-SEM) dataset (3 nm × 3 nm × 30 nm) [13] for comparison. The neuronal membrane structures in the PIB-SEM volume exhibit better continuity and are more easily identifiable across all three views compared to the ATUM-SEM volume. In both the XZ and YZ views, synaptic vesicles are clearly distinguishable in the PIB-SEM volume, while they appear blurry in the ATUM-SEM volume (Supplementary Fig. 5a). We quantify z-axial continuity using the Chunked Pearson Correlation Coefficient (CPC) [14] and the Structural Similarity Index Measure (SSIM) [15] (see *methods* for CPC and SSIM as continuity indicators). The PIB-SEM volume shows a 48.2% improvement in CPC and a 76.3% improvement in SSIM over the ATUM-SEM volume (Supplementary Fig. 5b), indicating better axial continuity.

As a result of superior axial resolution volumes generated by PIB-SEM, more efficient and accurate automated neuron segmentation could be achieved. The remarkable structural continuity across sections of the PIB-SEM volume facilitates seamless integration with existing general-purpose segmentation models, enabling the formation of an efficient human-in-the-loop segmentation pipeline with minimal human proofreading effort (Fig. 2a). Initially, we predict the membrane of the PIB-SEM volume using the original SegNeuron model [16], which demonstrates exceptional zero-shot performance. Experts then refine the precomputed results to establish the ground truth for model fine-tuning. Using the fine-tuned model, we performed watershed-based dense reconstruction of a 10 µm × 10 µm × 10 µm sub-volume (Supplementary Fig. 6). The finest substructures (dendritic spines and axons) can be well reconstructed (Fig. 2b), ensuring the applicability of connectomics. The superior axial resolution of PIB-SEM volume also enables more reliable connectome reconstruction. In contrast to the full resolution, the volume down-sampled to 40 nm along axial direction exhibits substantial merge or split errors, affecting the ultra-structures like spine necks and axons (Fig. 2c). These errors necessitate extensive manual proofreading, significantly increasing the labor costs for accurate connectome acquisition. These results demonstrate that the enhanced z-axis resolution of PIB preserves critical structural information essential for automated methods to achieve reliable segmentation performance.

**Fig. 2.**
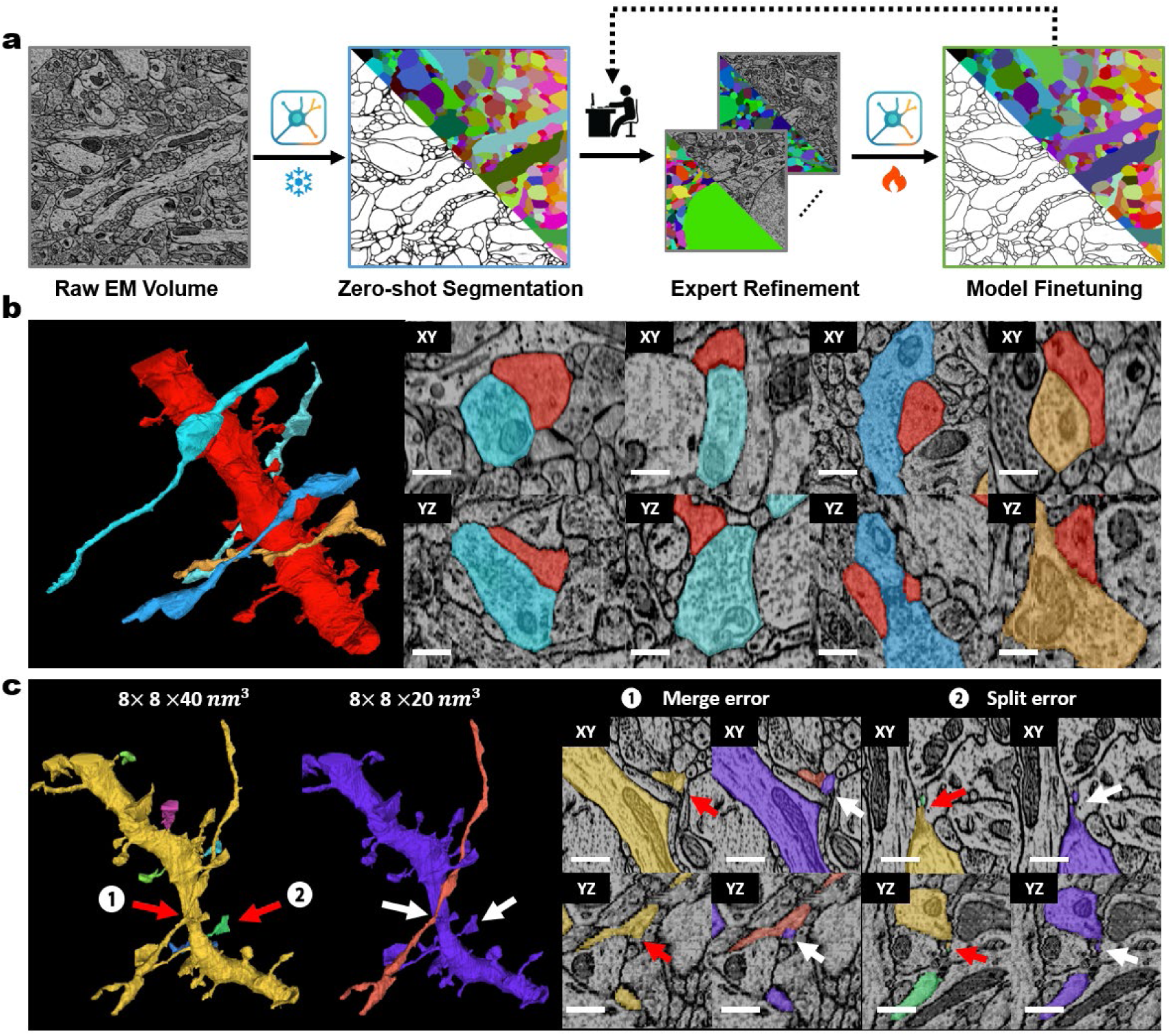
Automated segmentation of PIB-SEM data. **a**. Customized neural reconstruction workflow. Owing to high axial resolution and clearly defined ultrastructure in PIB-SEM volume, the general neuron segmentation model (SegNeuron) demonstrates strong reconstruction performance. Human experts only make connectivity corrections to the rough segmentation results, which are then used to fine-tune the model, further improving reconstruction accuracy. **b**. Local visualization of the neurite region, showing the connectivity between dendritic shafts and dendritic spines (scale bar: 500 nm). **c**. Comparison of segmentation model performance on the volumes with different axial resolution. The general model is fine-tuned and predicted on 40 nm and 20 nm axial resolution data. The results show that the 40 nm data introduce a large number of splitting and merging errors in finely connected regions. In contrast, the 20 nm data, thanks to its excellent spatial continuity, exhibits more accurate structural connectivity. The image on the right shows the visualization of the segmentation results on the original image (scale bar: 750 nm).

To demonstrate the applicability of PIB-SEM for millimeter-size samples, we section the rat cortex (see *Methods* for cortex of rat spanning all cortical layers), collect 150 serial sections with 100 nm thickness (Supplementary Fig. 7a). Five PIB-SEM cycles (10 × 10 × 20 nm^3^ voxels) are performed (Fig. 1h, Supplementary Fig. 7b). A volume of 1721 μm × 550 μm × 15 μm is reconstructed (Fig. 1h, Supplementary Fig. 7d), which spans all cortical layers (L1–L6) of the rat cerebral cortex. We can see the processing uniformity along the cortical layers (Supplementary Video 2).

To examine scalability toward large-area datasets required for neural circuit reconstruction, we extended PIB-SEM to a large contiguous portion of the mouse brain (see *Methods* for Coronal mouse brain spanning cortex to hypothalamus). Six consecutive sections (∼500 nm thickness each) were obtained from a mouse brain in the coronal plane containing cerebral cortex and hypothalamus regions, followed by ∼25 PIB-SEM acquisition cycles at 20 × 20 × 20 nm³ voxel resolution (Fig. 1h, Supplementary Fig. 8e). The resulting dataset covered 6470 μm × 3820 μm × 3 μm (Fig. 1h, Supplementary Fig. 8g). Due to the substantially increased section area, we used thicker sections (∼500 nm) to preserve structural integrity. Uniform sectioning by ultramicrotome across such ultralarge areas is technically challenging, resulting in uneven section thickness, with some regions slightly thicker or thinner than the nominal value. Etching rate also slightly varied, with regions near the sample center showing faster material removal, consistent with reduced staining penetration toward the interior and suggesting that staining gradients may affect local removal rates.

## Discussion

In summary, compared with existing techniques, PIB offers a unique complement to the field. FIB-SEM and GCIB provide nanometer-scale isotropic resolution but is constrained by sequential imaging, while ATUM-SEM enables large volumes but faces distortion and alignment issues. PIB combines the strengths of both, retaining ATUM’s wafer-based scalability while using ion-beam etching to achieve high axial resolution.

The three volumetric datasets acquired using PIB-SEM method demonstrate the significant potential of this technique for large-scale connectomics. By enabling simultaneous processing of hundreds of millimeter-scale sections with large-area thinning, high Z-accuracy, and complete retention of sections after thinning, this approach achieves both high throughput and reproducibility, significantly improving automated segmentation performance and reducing manual proofreading.

Thinning uniformity was evaluated on 4-inch wafers, with thickness deviations of less than 0.17 nm at the wafer center and less than 4.29 nm at the wafer edge. PIB can operate independently or in conjunction with multiple multibeam SEMs, significantly increasing the speed of large-volume imaging. The use of thicker sections also increases the reliability of serial section acquisition.

Nevertheless, PIB-SEM still has limitations. Section thickness varies due to mechanical deviations of ultramicrotome and the dullness of the diamond knife, particularly at the start of the section, leading to non-uniform thinning cycles. Future improvements in real-time thickness monitoring will help realize the full potential of PIB-SEM for whole-brain reconstruction.

Overall, PIB offers a parallelized, scalable, and high-fidelity imaging framework that expands the methodological landscape for connectomics and offers a possible approach for whole mouse brain neural circuit reconstruction.

## Methods

### Animals

All animal experiments were conducted in accordance with the regulations of the animal care agency and the Beijing Municipal Regulations for the management of laboratory animals. The rats used in this experiment were housed in the Experimental Animal Center of Tsinghua University, adhering to the standards for experimental rat care. The facility is graded as specific pathogen-free (SPF) to ensure that the animals are in a controlled environment. The housing conditions included a 12-hour light/dark cycle, a stable temperature of 24±1°C, relative humidity of 35±4%, with water and food available ad libitum.

### Samples preparation

All experimental protocols complied with relevant ethical regulations.

For the whole brain of mouse, the animals were anesthetized with isoflurane inhalation, they underwent cardiac perfusion fixation. First, the blood was quickly flushed with 0.1 M phosphate buffer (PB). Then, a solution of 2% paraformaldehyde (PFA) and 1.25% glutaraldehyde (GA) in 0.1 M PB was quickly perfused. The perfusion of the fixative was then slowed down and continued for no longer than 20 minutes. After the perfusion, the mouse’s brain were removed and then were fixed in a solution of 4% PFA and 2.5% GA in 0.1 M PB and kept at 4°C for 3 days. The samples were then washed using 0.1 M PB buffer. The post-fixation was carried out with 2% OsO₄ (Ted Pella, 18451), prepared in 0.1 M PB, for a duration of 9 days. Next, the samples were placed in a solution of 3% ferrocyanide (Sigma, 234125) in 0.1 M PB and incubated at room temperature for 6 days. Then, after washing three times with deionized water, the samples were incubated in 2% OsO₄ (Ted Pella, 18451) prepared in 0.1 M PB for 6 days. After that, the samples were thoroughly washed with deionized water and then incubated wiht freshly prepared pyrogallol solution (4%, in ddH_2_O) for 5 days. The samples were washed with deionized water followed by transfering the sample into a new tube containing freshly prepared 2% OsO_4_ aqueous solution for 6 days. After washing, they were incubated with 2% uranyl acetate (SPI-chem, 02624-AB) at 4°C for 2 days. Then, the samples were dehydrated through a graded ethanol (50%, 70%, 80%, 90%, 100%, 10 min each) and pure acetone on a horizontal shaker. Subsequently, the samples were infiltrated with Epon-812 resin (SPI, Epon-812), also on a horizontal shaker. The samples were immersed in epoxy/acetone solutions with increasing epoxy concentrations: 1:1, 2:1 and 3:1, and each immersion lasted 12-24 hours. Then, pure epoxy resin was infiltrated twice under negative pressure, each immersion lasted 24 hours. The last step was polymerization. The samples were placed in an embedding plate and processed at 60° C for 24 hours.

For the cortex of the rat brain, the rats were anesthetized with isoflurane inhalation followed cardiac perfusion fixation. After perfusion, the cortex region of the rat’s brain were removed and then were fixed in 4% PFA and 2.5% GA in 0.1 M PB for 2 hours. After washing, the samples were post-fixed with 2% OsO₄ and 3% ferrocyanide (Sigma, 234125) in 0.1 M PB for 2 hours. Then, after washing three times with deionized water, the samples were incubated in a solution of thiocarbohydrazide (TCH, Sigma, 223220) at 40°C for 45 min. After TCH treatment, the samples were thoroughly washed with deionized water and then incubated with 2% OsO_4_ for 90 min. The samples were then transferred to a new centrifuge tube for washing. After that, they were incubated with 1% uranyl acetate (SPI-chem, 02624-AB) at 4°C overnight. The samples dehydration was accomplished by transferring the samples through a graded series of acetone (50%, 70%, 80%, 90%, 100%) for 10 minutes each. For the sample embedding, a mixture of resin monomer and acetone was used to infiltrate the samples for 2 days in total. Finally, the resin-infiltrated samples were baked in an embedding oven at 60°C for 16 hours.

### Working principle of ion source

Unlike FIB and GCIB, PIB can generate a large-area uniform ion beam, thanks to the unique triple-grid regulation mechanism of the ion source (Fig. 1b, c, d; Supplementary Fig. 1d). First, in the plasma chamber of the ion source, high-energy electrons are excited by radio frequency (RF) and collided with argon gas molecules, causing ionization of the argon molecules and generating a large number of free electrons and Ar+ ions, thereby forming plasma. Then, after applying a voltage (80V) on the screen grid, Ar+ ions are extracted from the plasma chamber, while electrons are shielded within the plasma chamber. This effectively reduces the interference of low-energy electrons on the ion beam, ensuring the purity of the ion beam. Next, the ion beam is shaped by passing through the acceleration grid (-400V), which improves the uniformity and collimation of the ion beam. The voltage of the acceleration grid needs to be precisely adjusted based on the screen grid voltage. When the acceleration grid voltage is too low, the ion beam will not be effectively collimated. Conversely, if the voltage is too high, the ion beam will be completely absorbed by the acceleration grid, preventing the ion beam from being emitted properly. Finally, the ion beam undergoes deceleration through the grounded grid, forming a low-energy ion beam. The coordination of the acceleration grid and grounded grid is mainly used for shaping the ion beam, without changing its energy, which is determined by the voltage applied to the screen grid (Supplementary Fig. 1c,d). Ultimately, the Ar+ ions are extracted and shaped by the triple-grid system, forming a parallel ion beam with a diameter of 150 mm. The ion source is purchased from Beijing SHL Company.

### Sectioning

First, the Si wafer is cut into 2 cm×3 cm Si pieces using a dicing saw and cleaned and hydrophilized using a plasma cleaner. Then, an appropriate amount of water is added to the diamond knife boat (Supplementary Fig. 3a), and a thin layer of a mixture of Pattex glue (Henkel, Düsseldorf, Germany) and xylene (1:500) is applied on the side of the diamond knife that contacts the water, allowing the sections to stick together and form ribbons, thus preventing the sections from scattering. We use a room temperature diamond knife (35° angle) on an ultramicrotome (LEICA EM UC7) to slice from the rat brain embedded in Pon812 (Supplementary Fig. 3a). These sections are collected onto multiple Si pieces (Supplementary Fig. 3b; Supplementary Fig. 7a).

### Prefrontal cortex of rat

The sample from prefrontal cortex of a rat was sectioned. 1018 sections with 100 nm thickness were collected on three Si pieces (Supplementary Fig. 3b). Then the Si pieces were processed in the electron irradiation device (Bio-eBeam, China) to improve the conductivity of biological sections. After five PIB-SEM cycles were performed, these sections were almost etched away (Supplementary Fig. 3c). The PIB parameters used were as follows: ion beam energy of 80V, ion beam current of 100 mA, acceleration voltage of 400 V, glancing angle of 5°, neutralizing beam current of 400 mA, neutralizing gas flow rate of 7 sccm, ion beam diameter of 150 mm, and etching rate of 0.0277 nm/s; each thinning process lasted 720 seconds, with an etching depth of 20 nm. The SEM (Zeiss, Gemini 300) parameters used were: acceleration voltage of 6 kV, stage bias voltage - 5KV, aBSD detector, pixel size of 4 nm, and dwell time of 300 ns.

### Cortex of rat spanning all cortical layers

The sample from cortex of a rat spanning all cortical layers was sectioned. 150 sections with 100 nm thickness and 2220 μm×2050 μm size were collected on five Si pieces (Supplementary Fig. 7a). Then the Si pieces were processed in the electron irradiation device (Bio-eBeam, China) to improve the conductivity of biological sections. After five PIB-SEM cycles were performed, these sections are almost etched away (Supplementary Fig. 7b). The PIB parameters used were as follows: ion beam energy of 80V, ion beam current of 100 mA, acceleration voltage of 400V, glancing angle of 5°, neutralizing beam current of 400 mA, neutralizing gas flow rate of 7 sccm, ion beam diameter of 150 mm, etching rate of 0.0277 nm/s; each thinning process lasted 720 seconds, with an etching depth of 20 nm. The SEM (Zeiss, Gemini 300) parameters used were: acceleration voltage of 6 kV, stage bias voltage -5KV, aBSD detector, pixel size of 10 nm, and dwell time of 300 ns.

### Coronal mouse brain spanning cortex to hypothalamus

The sample from mouse brain were sectioned along the coronal plane, covering cerebral cortex and hypothalamus regions. Six sections, each ∼500 nm thick and measuring 6470 μm×3820 μm, were collected on a Si piece (Supplementary Figure 8a). Then the Si piece was processed in the electron irradiation device (Bio-eBeam, China) to improve the conductivity of biological sections. After ∼25 PIB-SEM cycles were performed, these sections are almost etched away (Supplementary Fig. 8e). The PIB parameters used were as follows: ion beam energy of 80V, ion beam current of 100 mA, acceleration voltage of 400V, glancing angle of 5 °, neutralizing beam current of 400 mA, neutralizing gas flow rate of 7 sccm, ion beam diameter of 150 mm, etching rate of 0.0277 nm/s; each thinning process lasted 560 seconds, with an etching depth of 20 nm. The SEM (Bio-eBeam, NavigatorSEM-100) parameters used were: acceleration voltage of 2 kV, BSD detector, pixel size of 20 nm, and dwell time of 25 ns. To reduce local variability in etching progression caused by uneven section thickness, we applied an inpainting-based correction method [17] to selected regions. This improved the continuity of the images while preserving the overall structural features. The correction was applied only for visualization purposes and did not affect global morphology or downstream quantitative analyses.

### Serial sections alignment

During the coarse alignment process, images of the stacks were down-sampled by 32X. We first performed pairwise alignment between every two adjacent images, extracted the corresponding points and applied affine transformations to estimate their deformations. In pairwise alignment, the sift method [18] was used to extract feature points of images, and the RANSAC [19] algorithm was used to filter corresponding point pairs. After pairwise alignment, we propagated the linear deformations across the entire stack [20]. In fine alignment process, we used an unsupervised optical flow network [21] to measure feature similarity between adjacent images. The optical flow network was then employed to estimate and compensate for cumulative registration error, thereby allowing for the reconstruction of the structure of biological tissues.

### Evaluation of thinning uniformity across 4-inch wafer

To assess the thinning uniformity of PIB across 4-inch wafer, five ribbons of consecutive sections with 100 nm thickness were collected on Si pieces separately. The Si pieces were mounted on the different positions of a 4-inch wafer using conductive tape, and labeled as Si1 to Si5(see Supplementary Fig. 2a). The first five sections in each ribbon were selected for thickness measurement.

The original thickness of each section was measured using Atomic Force Microscopy (AFM) and recorded in Supplementary Table 1, which varied from 80 nm to 110 nm. These sections were then processed in the electron irradiation device to improve the conductivity. After that, the thickness of each section was measured again using AFM (Supplementary Table 1). It was worth noting that electron beam irradiation, due to the negligible mass of electrons, did not produce milling effects but resulted in a thickness reduction of approximately 20 % to 30 %.

All of sections in 4-inch wafer underwent three thinning processes with 20 nm thickness. After each thinning process, the section thickness was measured and the thickness of removed sample was recorded in Supplementary Table 2. Considering the reduction of section thickness caused by electron irradiation step, we calculated the compensated thickness *ac_ibe* for removed sample by

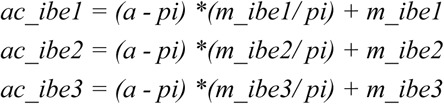

where *a* and *pi* were original and post-irradiation section thickness in Supplementary Table 1, *m_ibe* was the thickness of removed sample in Supplementary Table 2, the result of *ac_ibe* was detailed in Supplementary Table 3.

The results in Supplementary Table 3 indicate that the averaged thickness of removed sample at the center of the wafer deviates from the preset value by less than 0.17 nm, while at the wafer edge the deviation is less than 4.29 nm (see Supplementary Fig. 2b).

### CPC and SSIM as continuity indicators

In order to evaluate the axial continuity of the volume data, we used chunked pearson correlation coefficient (CPC) [14] and structural similarity index measure (SSIM) [15] as evaluation indicators. CPC is used to evaluate the correlation between two images in the local (block) area, while SSIM is used to measure the structural similarity of the two images. The aligned images in the datasets are evenly divided into image chunks without overlapping, as performed in [14]. The image chunks at the same horizontal position in adjacent sections are defined as matching pairs. The resolution of the input image was down-sampled to approximately 30 × 30 *nm*^2^/*pixel*, and each image chunk has a size of 64 × 64, which corresponds to a region of approximately 2 × 2 µ*m*^2^. Following the chunk division process, these criteria were computed for each matching pair independently.

The CPC and SSIM values (Supplementary Fig. 5b) of the PIB-SEM dataset are mainly concentrated in the high-value range (CPC > 0.64, SSIM > 0.57), indicating good continuity along axial direction. The CPC values of the ATUM-SEM dataset are concentrated around 0.468, which is significantly lower. The distribution of SSIM criteria also exhibits a similar trend.

### 3D reconstruction of the neurons in the cytoplasm

We applied a watershed-agglomeration method to obtain the dense reconstructions from PIB-SEM volumes. To construct the ground-truth dataset more efficiently, we annotated the neurons based on a precomputed segmentation instead of starting from scratch. Specifically, we first generated an initial segmentation of a 2000 × 2000 × 30 voxels PIB-SEM volume using a general-purpose segmentation model [SegNeuron], and then annotators checked through all the neurons and proofread the segmentation errors. The manually corrected blocks were subsequently utilized to fine-tune SegNeuron for learning high-quality affinity maps tailored to the target dataset. We employed the AdamW optimizer with a learning rate of 0.0001 to ensure stable convergence. The supervised training phase comprised 100,000 iterations with a batch size of 2. All models were implemented in PyTorch and executed on a single NVIDIA V100 GPU. Using the trained network, we ran the inference on a large volume with a size of 10 µm × 10 µm × 10 µm and obtained an affinity map of the same size. The 2D distance-transform-watershed algorithm was applied to each section to generate 2D superpixels. Then superpixels were agglomerated based on the multicut pipeline. The edge probability was computed by the mean affinity value accumulated on each edge. The hyper-parameter Beta was set as 0.25. We used GAEC to solve the multicut problem. For comparison, we downsampled the data in the z-direction to a voxel size of 8×8×40 nm^3 using nearest-neighbor interpolation, and applied the same training and inference pipeline.

---

**Supplementary Fig. 1.**
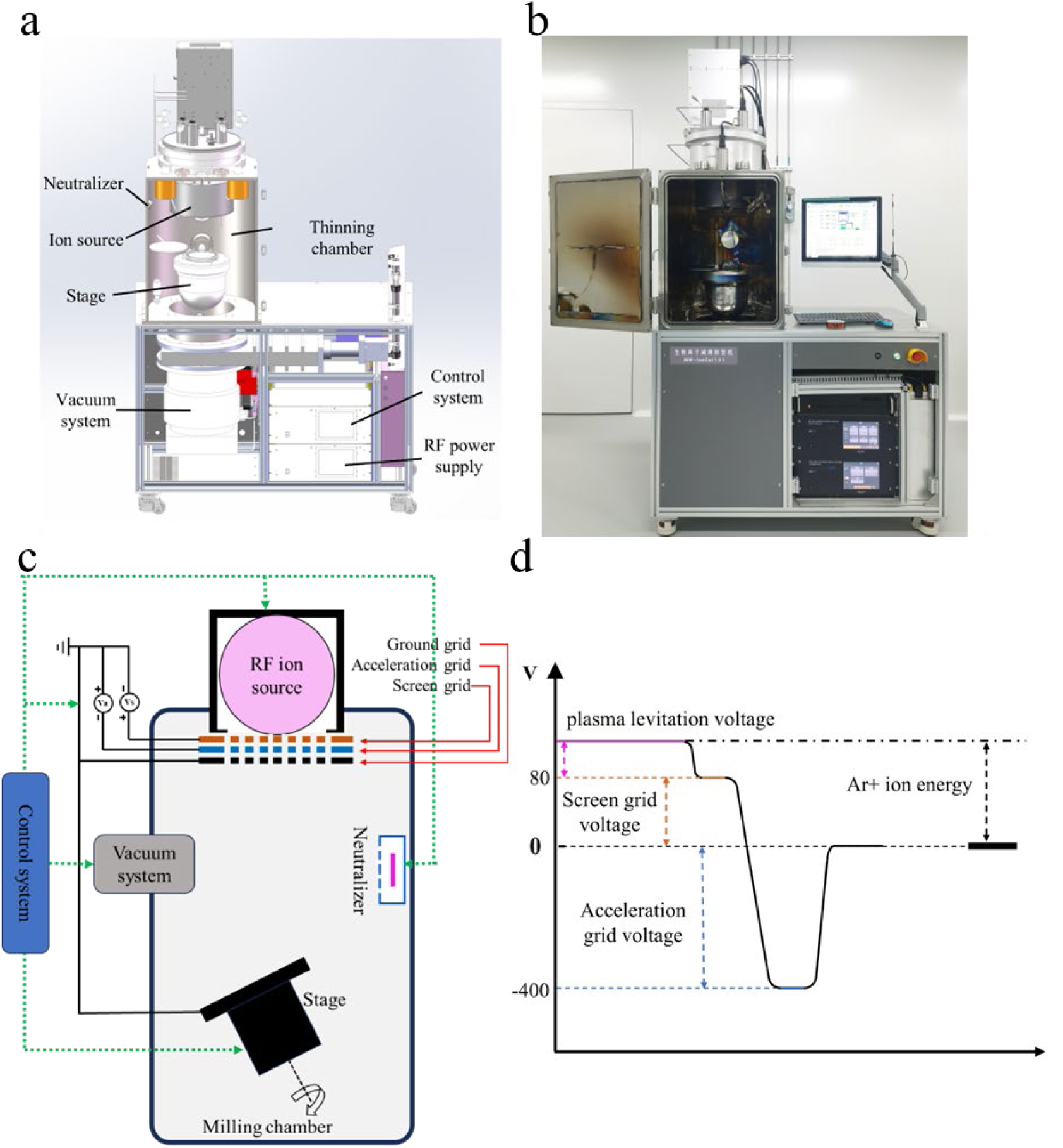
PIB prototype system. **a b,** 3D model and actual appearance of the PIB prototype. **c,** Operational mechanism of the PIB prototype. The parallel ion beam is emitted from the ion source. During the ion beam thinning process, to prevent the accumulation of positive charge (such as Ar⁺ ions) on the sample surface, which could affect the uniformity of the thinning, an equal number of electrons are released through the neutralizer to neutralize the surface charge in real-time, ensuring the stability of the thinning process. The control system is responsible for coordinating the operation of each module. **d,** Electric potential diagram of three-grid structure. In the plasma chamber of the ion source, high-energy electrons collide with argon gas molecules through radio frequency (RF) excitation, causing ionization of the argon molecules and generating a large number of free electrons and Ar+ ions, thereby forming a plasma. Under the negative voltage (80V) applied to the screen grid, the Ar+ ions are extracted from the plasma chamber, while the electrons are shielded within the plasma, reducing interference from low-energy electrons in the ion beam and ensuring the purity of the ion beam. The ion beam is then shaped by an acceleration grid (400V) to achieve high uniformity and collimation. The coordination between the acceleration grid and the ground grid is only used for shaping the ion beam and does not alter its energy.

**Supplementary Fig. 2.**
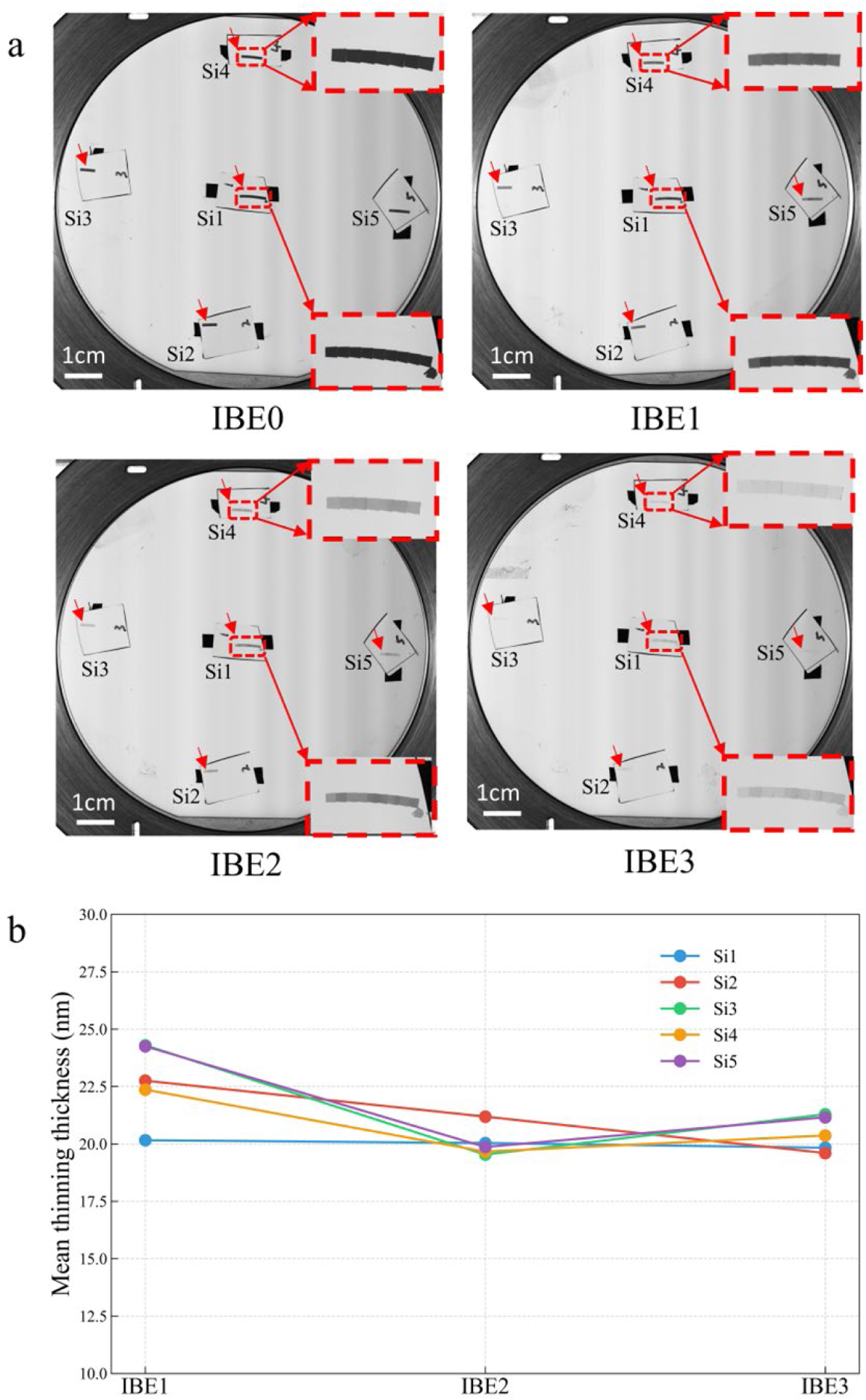
Evaluation of PIB thinning uniformity across 4-inch Si wafer. **a,** Five Si pieces (labeled Si1 to Si5) are mounted on the different positions of a 4-inch wafer. Each Si piece contains 5–9 sections, which are marked by red arrows. IBE0 shows the wafer before thinning. IBE 1, IBE2, and IBE3 correspond to the wafer after the first, second, and third PIB thinning, respectively. The thicknesses of the first five sections (arranged from right to left) on each Si piece are measured with AFM. **b**, the mean of the compensated thickness for removed sample on the Si pieces after each thing process.

**Supplementary Fig. 3.**
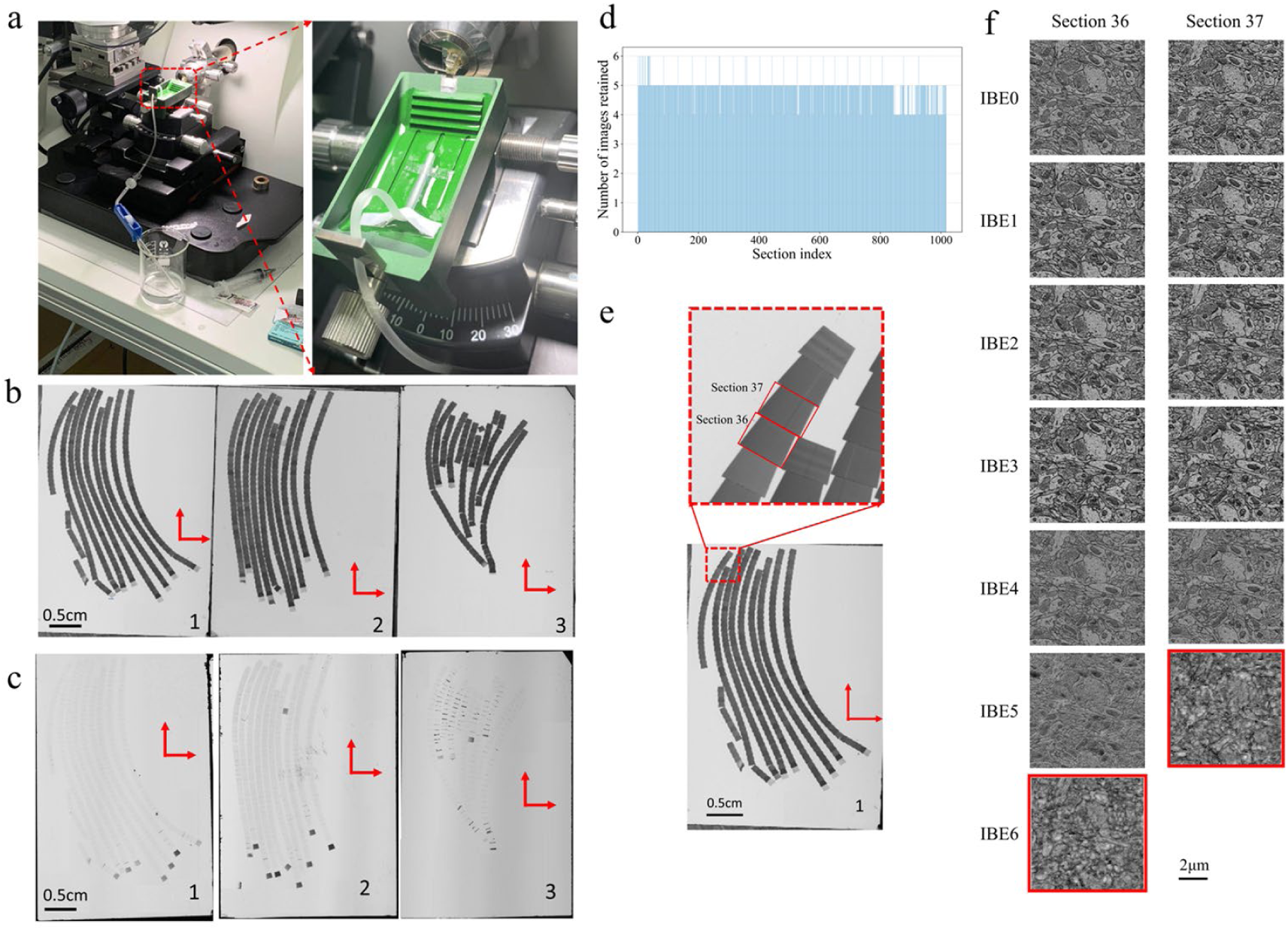
Serial sections collection and imaging. **a**, A custom-built device used for mounting ribbons of serial sections onto hydrophilized Si pieces. **b**, Images of Si pieces before PIB thinning. The sample from prefrontal cortex of a rat is sectioned using an ultramicrotome, resulting in 1018 section with 100 nm thickness. The sections are collected onto three Si pieces. **c**, Image of Si pieces after five PIB thinning process. **d**, Number of images retained for each of the 1018 sections after complete PIB thinning. **e**, Section 36 and 37 on the Si piece. **f**, SEM images of two consecutive sections (36 and 37) after multiple PIB thinning steps (20 nm thinning each). The red-boxed image shows that no meaningful content was left after complete thinning.

**Supplementary Fig. 4.**
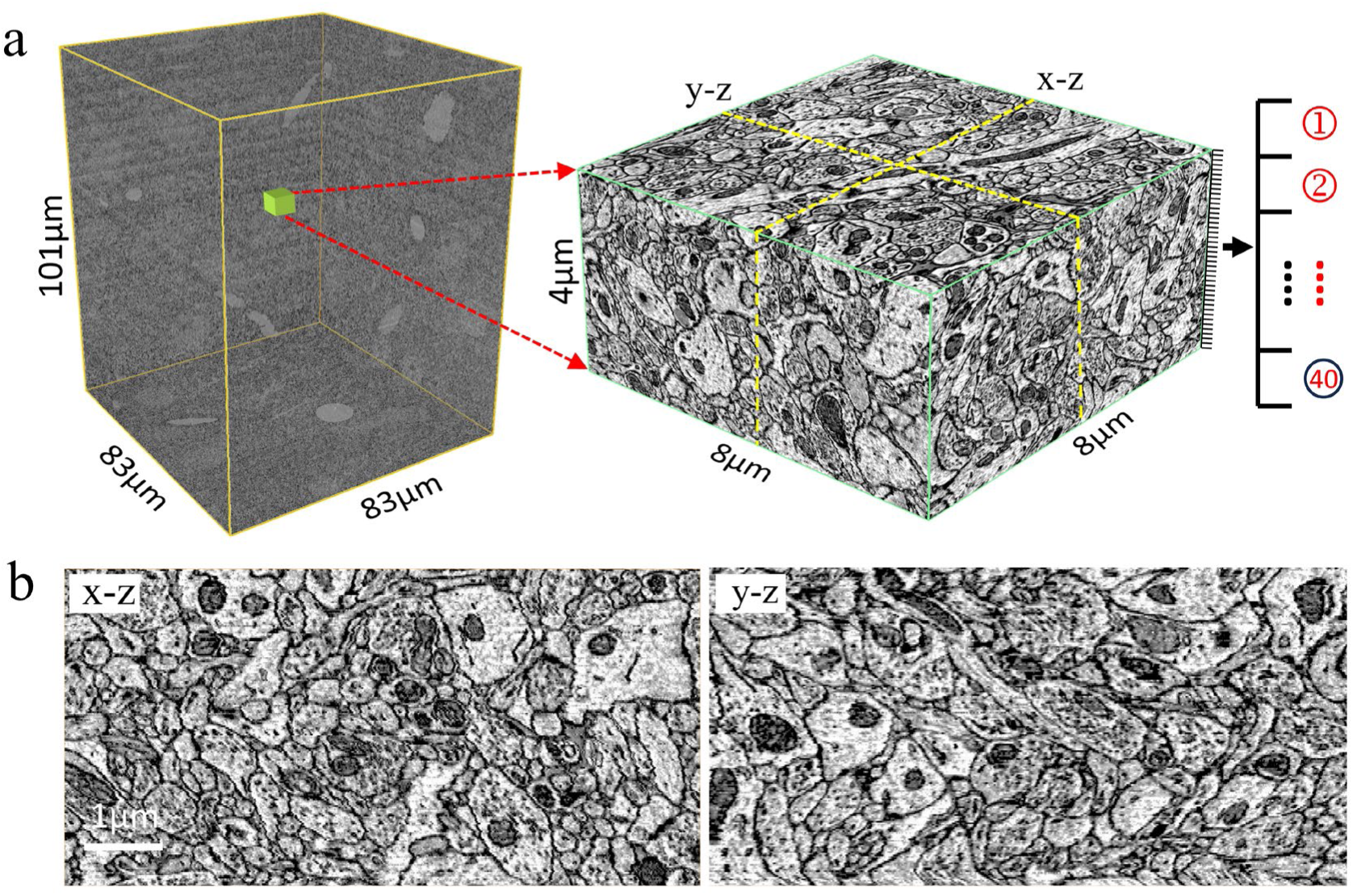
Image volume of prefrontal cortex of a rat. **a**, PIB-SEM volume (83 μm × 83 μm × 101 μm) spanning 1018 consecutive sections, and a sub-volume (8μm × 8μm × 4μm) spanning 40 consecutive sections. **b**, The x-z and y-z views of the yellow dashed line position in the sub-volume.

**Supplementary Fig. 5.**
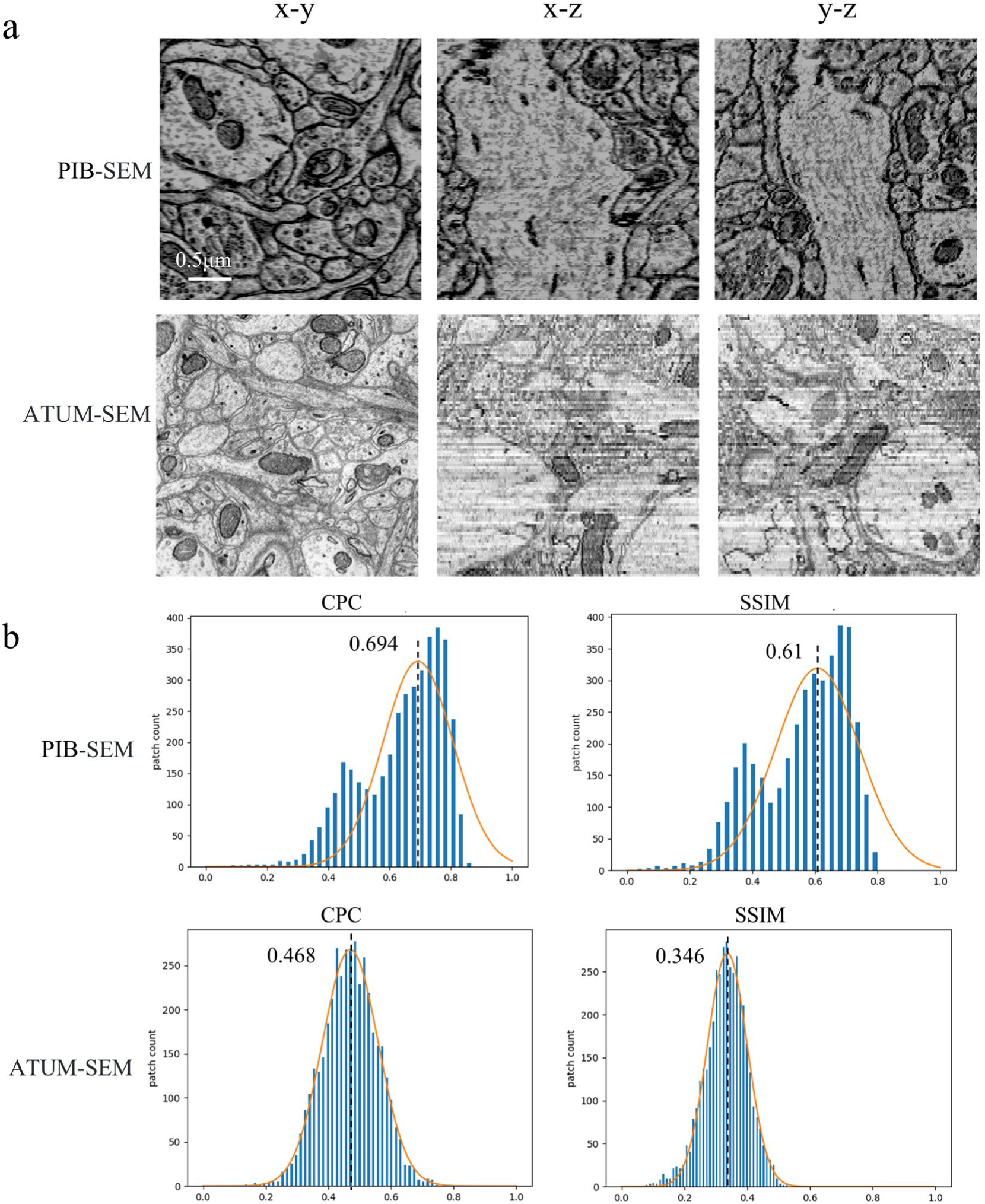
Comparison of different imaging volumes between the PIB-SEM volume (4 × 4 × 20 nm³ voxel) and the ATUM-SEM volume (3 × 3 × 30 nm³ voxel). **a**, Qualitative comparison of x-y, x-z, and y-z views. Compared to the ATUM-SEM volume, the neuronal membrane structures in the PIB-SEM volumes exhibit better continuity and are more easily identifiable across all three views. In the x-z and y-z views, synaptic vesicles are distinguishable in the PIB-SEM volume but blurred in the ATUM-SEM volume. Scale bar: 0.5 μm. **b**, Evaluation of z-axial continuity using CPC and SSIM. The brown line represents the Gaussian distribution fitting curve of the histogram. The value indicates the peak coordinate of the Gaussian distribution.

**Supplementary Fig. 6.**
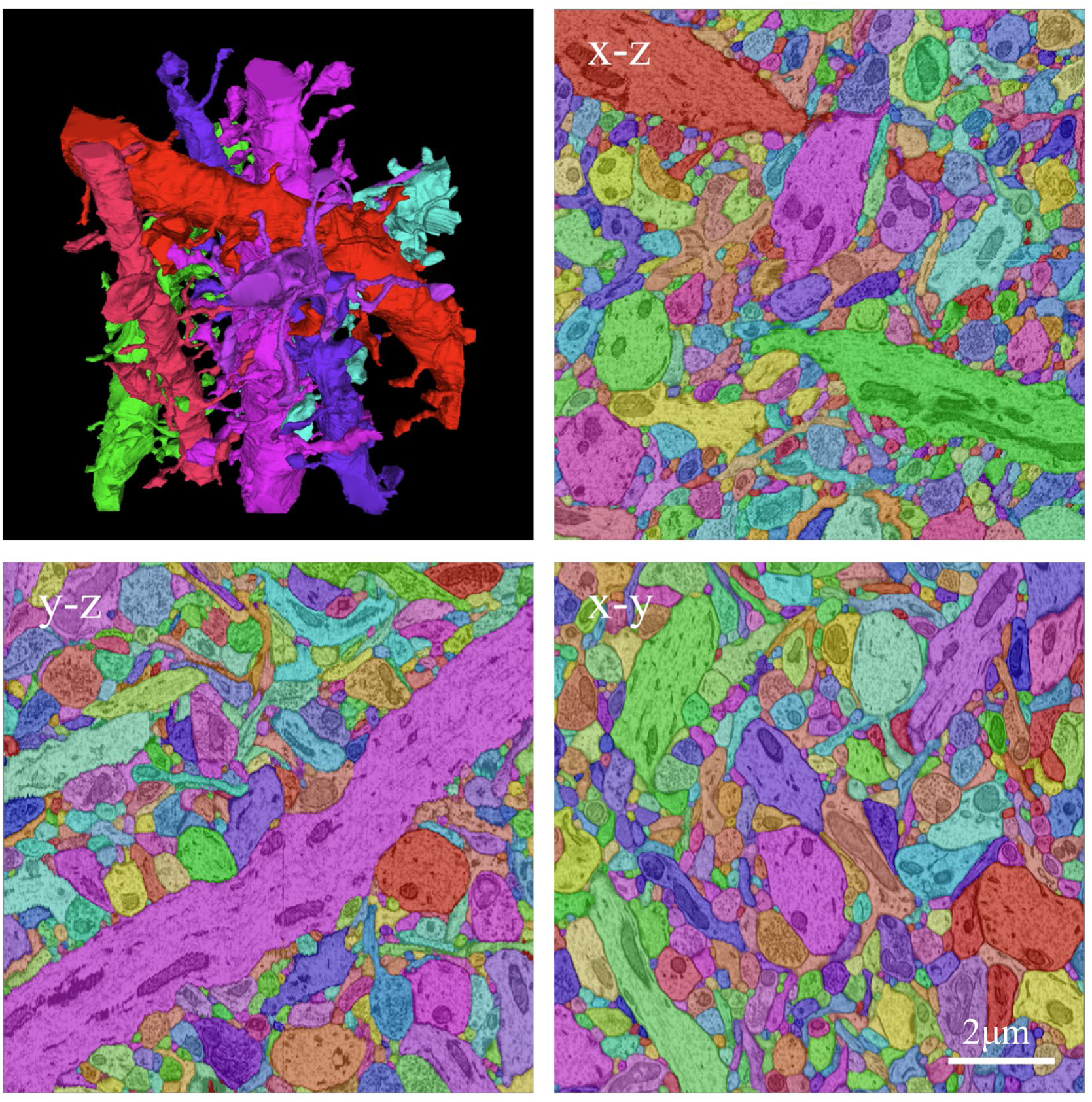
3D segmentation of a PIB-SEM volume. Dendritic reconstruction of the PIB-SEM volume (10 μm × 10 μm × 10 μm) and the x-z, x-y, and y-z views of the volume segmentation results.

**Supplementary Fig. 7.**
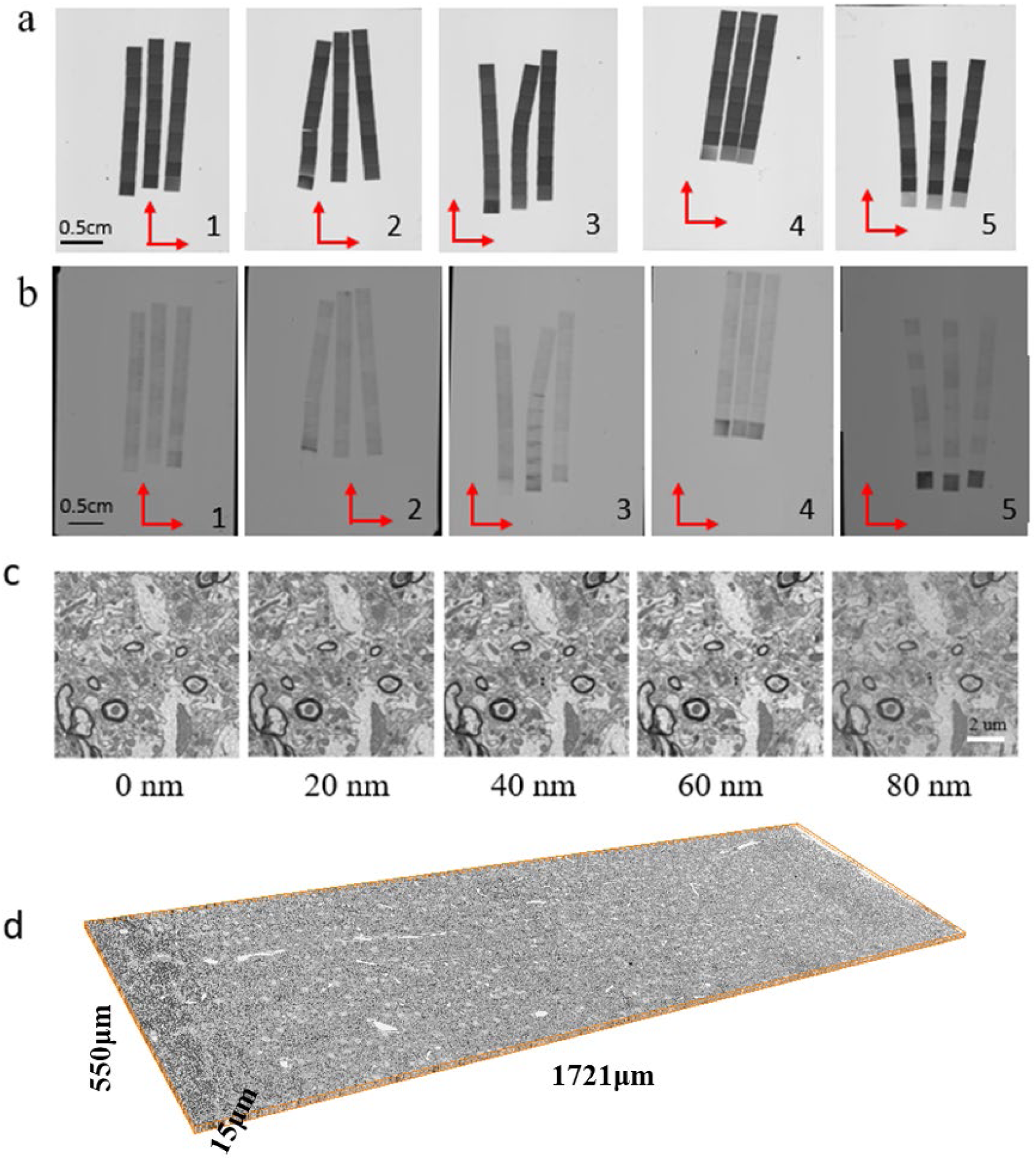
Image volume of cortex of rat spanning all cortical layers. **a,** 150 serial sections with 100 nm thickness are collected on five Si pieces. **b**, Image of Si pieces after five PIB thinning process. **c,** SEM images of a section after each PIB thinning process. **d,** Stitched stack of 1721 μm × 550 μm × 15 μm PIB-SEM volume.

**Supplementary Fig. 8.**
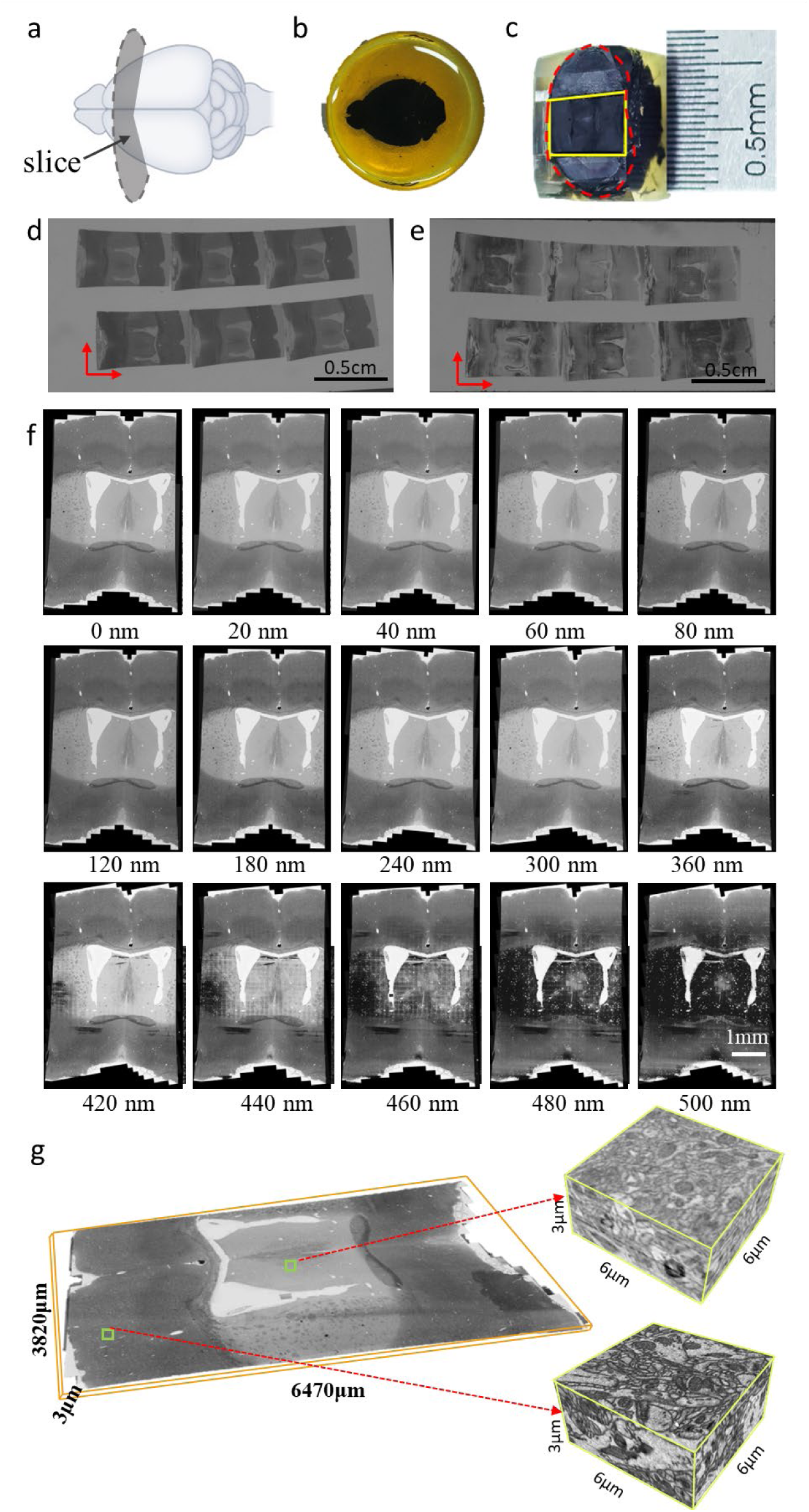
Image volume of coronal mouse brain spanning cortex to hypothalamus. **a,** Schematic diagram of the whole brain sample and section locations. **b,** whole brain sample **c,** Image of the sample showing the selected region of interest. The red outline indicates the sample contour, and the yellow region denotes the selected region of interest. **d,** 6 serial sections with ∼500 nm thickness are collected on a Si piece. **e**, Image of Si piece after ∼25 PIB thinning process. **f,** SEM images of a section after PIB thinning process. **g,** Stitched stack of 6470 μm × 3820 μm × 3 μm PIB-SEM volume.

**Supplementary Table 1.** Thickness (nm) of sections on each Si piece before and after electron irradiation.

|  | Si num | Section1<br>thickness | Section2<br>thickness | Section3<br>thickness | Section4<br>thickness | Section5<br>thickness |
| --- | --- | --- | --- | --- | --- | --- |
| Original<br>(*a*) | Si1 | 108.12 | 108.76 | 102.43 | 115.24 | 109.21 |
|  | Si2 | 92.98 | 91.30 | 95.66 | 91.51 | 85.26 |
|  | Si3 | 82.80 | 80.45 | 82.25 | 84.13 | 81.05 |
|  | Si4 | 91.78 | 97.79 | 91.70 | 99.86 | 98.15 |
|  | Si5 | 84.96 | 87.78 | 89.00 | 91.87 | 96.17 |
| post-Irradiation<br>(*pi*) | Si1 | 76.38 | 75.61 | 71.12 | 82.42 | 76.91 |
|  | Si2 | 69.93 | 69.94 | 70.11 | 68.86 | 64.25 |
|  | Si3 | 60.22 | 56.78 | 58.99 | 61.63 | 58.00 |
|  | Si4 | 67.20 | 68.81 | 66.48 | 70.71 | 69.21 |
|  | Si5 | 64.69 | 67.10 | 68.53 | 70.45 | 74.34 |

**Supplementary Table 2.** Thickness (nm) of removed sample after each PIB thinning.

|  | Si num | Section1<br>thickness | Section2<br>thickness | Section3<br>thickness | Section4<br>thickness | Section5<br>thickness |
| --- | --- | --- | --- | --- | --- | --- |
| IBE1<br>(*m_ibe1*) | Si1 | 14.74 | 13.47 | 14.38 | 15.39 | 12.90 |
|  | Si2 | 17.93 | 15.52 | 16.11 | 17.76 | 18.14 |
|  | Si3 | 16.06 | 17.36 | 16.36 | 19.29 | 18.38 |
|  | Si4 | 16.01 | 14.26 | 13.85 | 19.13 | 16.61 |
|  | Si5 | 19.65 | 18.08 | 18.8 | 17.21 | 19.29 |
| IBE2<br>(*m_ibe2*) | Si1 | 14.17 | 14.52 | 12.91 | 14.32 | 14.51 |
|  | Si2 | 14.50 | 17.62 | 14.98 | 14.45 | 18.11 |
|  | Si3 | 14.29 | 13.92 | 14.33 | 11.88 | 15.81 |
|  | Si4 | 14.06 | 12.07 | 16.53 | 13.78 | 13.88 |
|  | Si5 | 14.55 | 13.85 | 15.20 | 16.53 | 16.08 |
| IBE3<br>(*m_ibe3*) | Si1 | 13.48 | 12.38 | 16.68 | 15.66 | 11.51 |
|  | Si2 | 13.98 | 14.47 | 17.71 | 15.62 | 11.75 |
|  | Si3 | 13.46 | 14.53 | 14.78 | 18.02 | 15.83 |
|  | Si4 | 14.07 | 17.16 | 13.54 | 15.26 | 12.69 |
|  | Si5 | 15.59 | 16.58 | 15.77 | 16.52 | 16.70 |

**Supplementary Table 3.** Compensated thickness (nm) of removed sample after each PIB thinning.

|  | Si num | Section1<br>thickness | Section2<br>thickness | Section3<br>thickness | Section4<br>thickness | Section5<br>thickness |
| --- | --- | --- | --- | --- | --- | --- |
| IBE1<br>(*ac_ibe1*) | Si1 | 20.86 | 19.37 | 20.72 | 21.53 | 18.32 |
|  | Si2 | 23.84 | 20.26 | 21.99 | 23.61 | 24.08 |
|  | Si3 | 22.08 | 24.59 | 22.8 | 26.32 | 25.68 |
|  | Si4 | 21.87 | 20.27 | 19.11 | 27.02 | 23.55 |
|  | Si5 | 25.81 | 23.65 | 24.41 | 22.44 | 24.96 |
| IBE2<br>(*ac_ibe2*) | Si1 | 20.06 | 20.89 | 18.60 | 20.03 | 20.61 |
|  | Si2 | 19.28 | 23.00 | 20.43 | 19.20 | 24.03 |
|  | Si3 | 19.64 | 19.73 | 19.98 | 16.22 | 22.10 |
|  | Si4 | 19.20 | 17.15 | 22.8 | 19.47 | 19.68 |
|  | Si5 | 19.11 | 18.12 | 19.73 | 21.55 | 20.81 |
| IBE3<br>(*ac_ibe3*) | Si1 | 19.08 | 17.81 | 24.03 | 21.90 | 16.35 |
|  | Si2 | 18.59 | 18.89 | 24.17 | 20.76 | 15.60 |
|  | Si3 | 18.51 | 20.59 | 20.61 | 24.60 | 22.12 |
|  | Si4 | 19.22 | 24.38 | 18.67 | 21.55 | 18.00 |
|  | Si5 | 20.48 | 21.69 | 20.48 | 21.54 | 21.60 |

